# Geometric Deep Learning of the Human Connectome Project Multimodal Cortical Parcellation

**DOI:** 10.1101/2021.08.18.456790

**Authors:** Logan Z. J. Williams, Abdulah Fawaz, Matthew F. Glasser, A. David Edwards, Emma C. Robinson

## Abstract

Understanding the topographic heterogeneity of cortical organisation is an essential step towards precision modelling of neuropsychiatric disorders. While many cortical parcellation schemes have been proposed, few attempt to model inter-subject variability. For those that do, most have been proposed for high-resolution research quality data, without exploration of how well they generalise to clinical quality scans. In this paper, we benchmark and ensemble four different geometric deep learning models on the task of learning the Human Connectome Project (HCP) multimodal cortical parcellation. We employ Monte Carlo dropout to investigate model uncertainty with a view to propagate these labels to new datasets. Models achieved an overall Dice overlap ratio of >0.85 ± 0.02. Regions with the highest mean and lowest variance included V1 and areas within the parietal lobe, and regions with the lowest mean and highest variance included areas within the medial frontal lobe, lateral occipital pole and insula. Qualitatively, our results suggest that more work is needed before geometric deep learning methods are capable of fully capturing atypical cortical topographies such as those seen in area 55b. However, information about topographic variability between participants was encoded in vertex-wise uncertainty maps, suggesting a potential avenue for projection of this multimodal parcellation to new datasets with limited functional MRI, such as the UK Biobank.

## Introduction

Cortical parcellation is the process of segmenting the cerebral cortex into functionally specialised regions. Most often, these are defined using sulcal morphology [5], and are propagated to individuals from a population-average template (or set of templates) based on the correspondence of cortical shape [28,17]. By contrast, while it is possible to capture subject-specific cortical topography from functional imaging in a data-driven way [14], it is difficult to perform populationbased comparisons with these approaches as they typically result in parcellations where the number and topography of the parcels vary significantly across subjects [15]. Notably, even following image registration methods that use both structural and functional information [26,25], considerable topographic variation remains across individuals [10,19].

Recently, [10] achieved state-of-the-art cortical parcellation through hand annotation of a group-average multimodal magnetic resonance imaging (MRI) atlas from the Human Connectome Project (HCP). Specifically, a sharp group average of cortical folding, cortical thickness, cortical myelination, task and resting state functional MRI (fMRI), were generated through novel multi-modal image registration [25] driven by ‘areal features’: specifically T1w/T2w ratio (cortical myelin) [13] and cortical fMRI; modalities which are known to more closely reflect the functional organisation of the brain. This improved alignment allowed for manual annotation of regional boundaries via identification of sharp image gradients, consistent across modalities. With this group average template, they trained a multi-layer perceptron (MLP) classifier to recognise the multimodal ‘fingerprint’ of each cortical area. This approach allowed [10] to propagate parcellations from labelled to unlabelled subjects, in a registration-independent manner, also providing an objective method to validate parcellation in an independent set of test participants. This classifier detected 96.6% of the cortical areas in test participants, and could correctly parcellate areas in individuals with atypical topography [10].

However, even in this state-of-the-art approach, the classifier was still unable to detect 3.4% of areas across all subjects [10]. Moreover, they were unable to replicate previously identified parcels in regions such as the orbitofrontal cortex [23] and the association visual cortex [1]. It is also unknown whether this classifier generalises to different populations with lower quality data, for example the UK Biobank [21] and the Developing Human Connectome Project [20]. Thus, development of new tools that improve upon areal detection and allow generalisation of this parcellation to new populations with less functional MRI data is warranted. To this end, we consider convolutional neural networks (CNNs), which have proven state-of-the-art for many 2D and 3D medical imaging tasks [16, 3]. More specifically, we benchmark a range of different geometric deep learning (gDL) frameworks, since these adapt CNNs to irregular domains such as surfaces, meshes and graphs [2].

The specific contributions of this paper are as follows:

1. We propose a novel framework for propagating the HCP cortical parcellation [10] to new surfaces using gDL methods. These offer a way to improve over vertex-wise classifiers (as used by [10]) by additionally learning the spatial context surrounding different image features
2. Since gDL remains an active area of research, with several *complementary* approaches for implementing surface convolutions, we explore the potential to improve performance by ensembling predictions made across a range of models.
3. Given the degree of heterogeneity and anticipated problems in generalising to new data, we return estimates of model uncertainty using techniques for Bayesian deep learning implemented using Monte Carlo dropout [8].

## Methods

### Participants and image acquisition

A total of 390 participants from the HCP were included in this study. Acquisition and minimal preprocessing pipelines are described in [12]. Briefly, modalities included T1w and T2w structural images, task-based and resting state-based fMRI images, acquired at high spatial and temporal resolution on a customized Siemens 3 Tesla (3T) scanner [12]. From these, a set of 110 features were derived and used as inputs for cortical parcellation: 1 thickness map corrected for curvature, 1 T1w/T2w map [13], 1 surface curvature map, 1 mean task-fMRI activation map, 20 task-fMRI component contrast maps, 77 surface resting state fMRI maps (from a d=137 independent component analysis), and 9 visuotopic features. This differs from 112 features used by the MLP classifier in [10] in that artefact features were not included and visuotopic spatial regressors were included. Individual subject parcellations *predicted* by the MLP classifier were used as labels for training each gDL model, as there are no ground truth labels available for multimodal parcellation in the HCP.

### Modelling the cortex as an icosphere

For all experiments, the cortical surface was modelled as a regularly tessellated icosphere: a choice which reflects strong evidence that, for many parts of the cortex, cortical shape is a poor correlate of cortical functional organisation [7, 10]. Icospheres also offer many advantages for deep learning. Since their vertices form regularly spaced hexagons, icospheric meshes allow consistently shaped spatial filters to be defined and lend themselves to straightforward upsampling and downsampling. This generates a hierarchy of regularly tessellated spheres over multiple resolutions, which is particularly useful as it allows deep learning models to aggregate information through pooling.

### Image processing and augmentation

Spherical meshes and cortical metric data (features and labels) for each subject were resampled from the 32k (FS_LR) HCP template space [30], to a sixth-order icosphere (with 40,962 vertices). Input spheres were augmented using non-linear spherical warps estimated by: first, randomly displacing the vertices of a 2nd order icospheric mesh; then propagating these deformations to the input meshes using barycentric interpolation. In total, 100 warps were simulated, and these were randomly sampled from during training. Cortical metric data were then normalised to a mean and standard deviation of 0 and 1 respectively, using precomputed group means and standard deviations per feature.

### Model Architecture & Implementation

Geometric convolutions may be broadly classified into spatial or spectral methods, which reference the domain that the convolution is computed in (see [2, 9] for more details). In brief, spatial methods [32, 22] simulate the familiar concept of passing a localised filter over the surface. In practice, while expressive, such methods often approximate mathematically correct convolutions; since, due to lack of a single, fixed coordinate system it is not possible to slide a filter over a curved surface whilst maintaining consistent filter orientation. Spectral methods, on the other hand, utilise an alternate representation in which the (generalised) Fourier transform of a convolution of two functions may be represented by the product of their Fourier transforms. As full spectral methods are computationally expensive, it is standard practice to address this through polynomial approximation [4].

Each method therefore results in different compromises, and for that reason offers complementary solutions, which in principle may be combined to improve performance. In this paper, we therefore benchmark and ensemble two spatial networks: *Spherical U-Net* [32] and *MoUNet* [22]; and two spectral (polynomial approximation) methods: *ChebNet* [4] and *GConvNet* [18].

In each case, methods were implemented with a U-Net [27] like architecture with a 6-layer encoder and decoder, and upsampling was performed using transpose convolution (as implemented by [32]). Code for Spherical U-Net was implemented from its GitHub repository^6^ and ChebNet, GConvNet and MoUnet were written using PyTorch Geometric [6]. Optimisation was performed using Adam, with an unweighted Dice loss and learning rates: 1 × 10^−3^ (for Spherical U-Net) and 1 × 10^−4^ (for all other models). All models were implemented on a Titan RTX 24GB GPU, with batch size limited to 1 due to memory constraints (resulting from the high dimension of input channels). Models were trained and tested with a train/validation/test split of 338/26/26, using data from both left and right hemispheres. Following training, an unweighted ensemble approach was taken, where one-hot encoded predictions for a single test subject were averaged across all gDL models. Model performance was also assessed using weighted recall and precision scores. Finally uncertainty estimation was implemented using test-time dropout [8] (with *p* = 0.2, the probability of an input channel being dropped). Vertex-wise uncertainty maps were produced by repeating dropout 200 times, and then calculating the standard deviation across each vertex of the predicted parcellation per subject.

## Results

Table 1 shows the overall performance each model on HCP parcellation using single subject cortical maps predicted by the HCP MLP classifier. All methods perform well, achieving a Dice overlap ratio of >0.85, recall score of >0.82 and precision score of >0.85. The mean and standard deviation Dice overlap ratio, recall and precision scores per area are shown for GConvNet (the best performing model) in Figure 1a-c. V1 and cortical areas in the parietal lobe had higher a mean and lower standard deviation Dice overlap ratio, whilst cortical areas in the medial frontal lobe, occipital pole and insula had lower a mean and higher standard deviation Dice overlap ratio. At the level of a single cortical region, mean and standard deviation Dice overlap ratio varied across models (figure 1b-d), and this regional variability was utilised through an ensemble approach to improve parcellation performance (table 1).

**Table 1.** Mean ± standard deviation Dice overlap ratio, recall, and precision for all four geometric deep learning methods and the unweighted ensemble approach

| Method | Dice overlap ratio | Recall | Precision |
| --- | --- | --- | --- |
| ChebNet | $0.871 \pm 0.021$ | $0.839 \pm 0.024$ | $0.862 \pm 0.016$ |
| **GConvNet** | **$0.875 \pm 0.020$** | **$0.843 \pm 0.230$** | **$0.865 \pm 0.015$** |
| MoUNet | $0.873 \pm 0.021$ | $0.841 \pm 0.023$ | $0.864 \pm 0.015$ |
| Spherical UNet | $0.860 \pm 0.021$ | $0.825 \pm 0.022$ | $0.851 \pm 0.013$ |
| **Ensemble** | **$0.880 \pm 0.019$** | **$0.848 \pm 0.022$** | **$0.860 \pm 0.019$** |

**Fig. 1.**
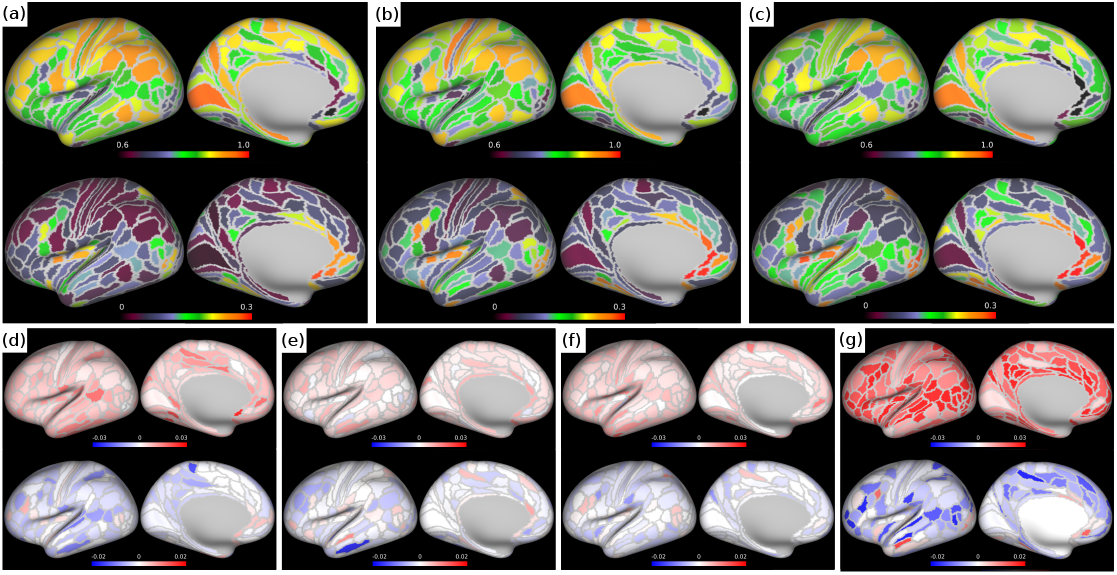
Mean (top row) and standard deviation (bottom row) (a) Dice overlap ratio, (b) recall score (c) and precision score per region for gDL ensemble. Mean (top row) and standard deviation (bottom row) Dice overlap ratio per region for (d) ensemble - ChebNet, (e) ensemble - GConvNet, (f) GConvNet - MoUNet, and (g) ensemble - Spherical UNet

The ability of gDL models to detect atypical cortical topography was assessed qualitatively in a test-set participant where area 55b was split into three distinct parcels by the frontal and posterior eye fields [11]. This showed that, while none of the gDL models predicted this split (Figure 2b), vertex-wise uncertainty maps highlighted the split as a region of uncertainty. Figure 3b demonstrates that the most likely labels for this subject (at the vertex marked with a white dot) were the frontal eye fields (184/200 epochs) and area 55b (16/200 epochs). By contrast, when compared to a similar vertex location in a subject with typical parcellation in area 55b (figure 3c), there was no uncertainty in the estimated label, predicting area 55b across all epochs.

**Fig. 2.**
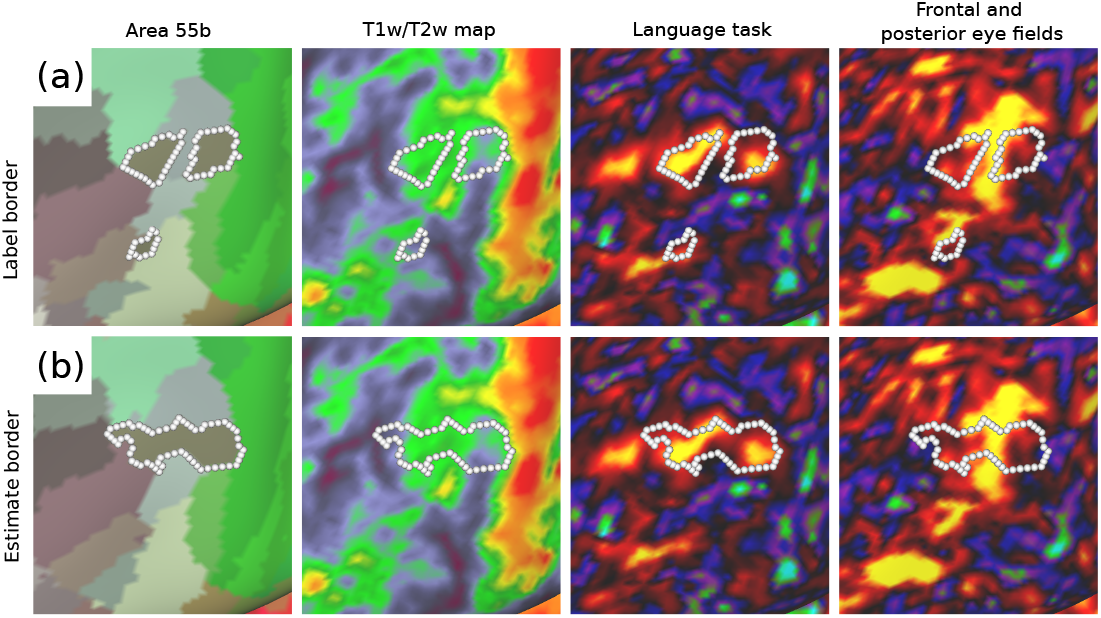
Label border (a) and estimated (b) border predicted by gDL ensemble for test set participant with atypical area 55b topography. Borders are overlaid on T1w/T2w map, and functional connectivity map from the HCP language task, and functional connectivity map highlighting the frontal and posterior eye fields.

**Fig. 3.**
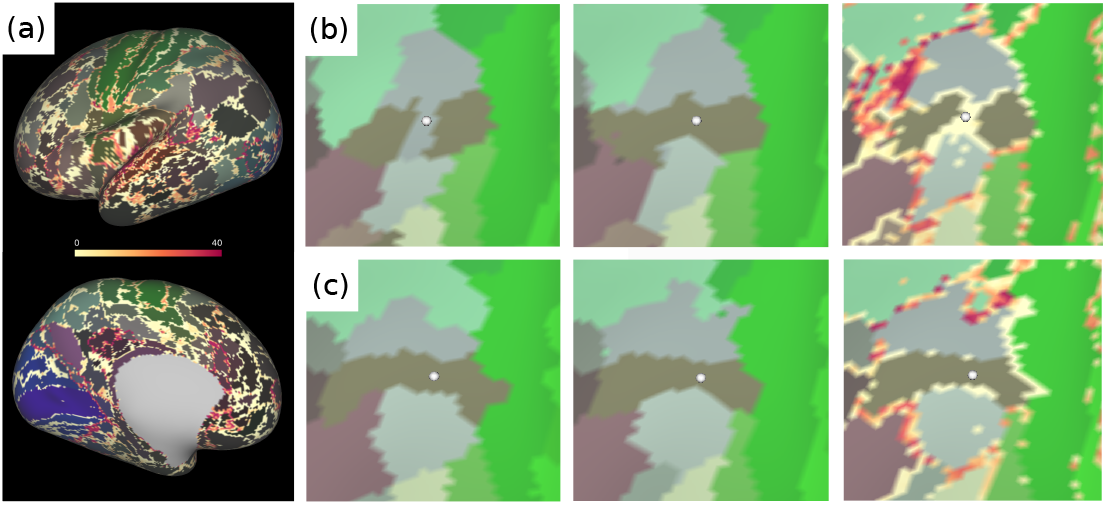
(a) Example of a vertex-wise uncertainty map produced using Monte Carlo dropout (MoUNet) (b) From left to right: label, estimate and vertex-wise uncertainty map in subject with *atypical* topography of area 55b. (c) from left to right: label, estimate and vertex-wise uncertainty map in subject with *typical* topography of area 55b.

Beyond area 55b, gDL models often predicted cortical areas as single contiguous parcels, whereas the HCP MLP classifier predicted some cortical areas as being comprised of several smaller, topographically-distinct parcels. This uncertainty relative to the MLP is further emphasised by the findings from the Monte Carlo dropout uncertainty modelling which showed that areas of uncertainty tended to be greatest along the boundaries between regions, and were higher in locations where >2 regions met.

## Discussion

Developing methods that capture the topographic variability of cortical organisation is essential for precision modelling of neuropsychiatric disorders. Here we show that gDL methods achieve good performance in predicting subject’s cortical organisation, when trained on labels output from the HCP MLP classifier.

Even though overall metrics of regional overlap were high, there was marked variability across cortical areas. These findings are in part a consequence using an unweighted Dice loss; since, in this case, mislabelling single vertices of smaller cortical areas will have less impact than for larger ones [24]. This is reflected in the results above, where larger regions e.g V1, and cortical areas in the parietal lobe, had higher mean and lower standard deviation Dice overlap ratio, recall and precision score per region. This is compared to smaller regions in the medial frontal lobe, insula, and lateral occipital lobe, which had lower mean and higher standard deviation. This inherent limitation of the Dice overlap ratio might also explain why GConvNet (the gDL method with the smallest kernel size) performed the best, as it was capable of learning very localised features. The variation in performance across cortical areas also differed between models, which suggests that each gDL model is learning a different set of features. This was expected given the theoretical differences in how in each model’s convolution is defined. Utilising these differences in an emsemble approach improved Dice overlap ratio by 0.005 (0.5%) above GConvNet, which translates to an overlap improvement of 200 vertices on a 6th-order icosphere (on the same icosphere, area 55b is only 123 vertices in size).

Although not described here, we also trained these gDL methods using a generalised (weighted) Dice overlap ratio as described by [29] that is designed to address class imbalance, but found that it did not perform as well as the unweighted Dice overlap ratio. This suggests that future work on improving model performance should, in part, address the limitations of common image segmentation losses in the context of multimodal cortical parcellation.

Qualitative assessment of gDL model performance on cortical parcellation is essential for investigating topographic variability, as this information is is not fully captured by performance metrics. Although subjects with atypical topography of area 55b were included in the training set, none of the gDL methods were able to correctly identify this topography in a test-set subject. Specifically, all models predicted area 55b as a contiguous parcel compared to the HCP MLP prediction where it was split into three smaller areas by the frontal and posterior eye fields. The atypical topography of area 55b in this subject was confirmed manually from the features known to contribute to its multimodal fingerprint (namely, T1w/T2w ratio, the HCP language task contrast “Story vs. Baseline” and resting-state functional connectivity map) [11].

The performance of the gDL models in area 55b highlights the overall tendency of these models to predict cortical areas as contiguous regions compared to those predicted by the HCP MLP classifier. This behaviour might be a result of CNNs learning spatial context, and a strong bias towards learning typical topographic organisation due to downsampling and skip connections in the U-Net architecutre. In contrast, the HCP MLP was trained to classify each vertex independently using limited spatial context (30mm radius searchlight across the surface) [10]. The importance of spatial contiguity in defining cortical areas is unknown, but given the lack of ground truth it is difficult to evaluate which approach is more accurate without extensive further qualitative and quantitative evaluation. However, these results do suggest each model introduces unique biases that need to be accounted for when investigating cortical organisation and neuropsychiatric disorders.

Achieving multimodal cortical parcellation in datasets beyond the HCP will be invaluable for precision modelling of neuropsychiatric disorders. However, generalising these multimodal labels to other datasets such as the UK Biobank (healthy ageing adults) [21] and the Developing Human Connectome Project (term and preterm neonates) [20] is challenging due to differences in population demographics and data acquisition (less and lower quality). The vertex-wise uncertainty maps introduced here provide a quantitative method to evaluate label propagation, which also could be used to inform post-processing of individual participant cortical parcellations, similar in nature to [10]. The information about topographic variability encoded in these vertex-wise maps might also provide a way to investigate atypical topography in less explored cortical areas.

## Acknowledgements

Data were provided by the Human Connectome Project, WU-Minn Consortium (Principal Investigators: David Van Essen and Kamil Ugurbil; 1U54MH091657) funded by the 16 NIH Institutes and Centers that support the NIH Blueprint for Neuroscience Research; and by the McDonnell Center for Systems Neuroscience at Washington University [31].

## Footnotes

6 https://github.com/zhaofenqiang/Spherical_U-Net

## References

1. Abdollahi, R.O., Kolster, H., Glasser, M.F., Robinson, E.C., Coalson, T.S., Dierker, D., Jenkinson, M., Van Essen, D.C., Orban, G.A.: Correspondences between retinotopic areas and myelin maps in human visual cortex. Neuroimage 99, 509–524 (2014)

2. Bronstein, M.M., Bruna, J., LeCun, Y., Szlam, A., Vandergheynst, P.: Geometric deep learning: going beyond euclidean data. IEEE Signal Processing Magazine 34(4), 18–42 (2017)

3. Chen, H., Dou, Q., Yu, L., Qin, J., Heng, P.A.: Voxresnet: Deep voxelwise residual networks for brain segmentation from 3d mr images. NeuroImage 170, 446–455 (2018)

4. Defferrard, M., Bresson, X., Vandergheynst, P.: Convolutional neural networks on graphs with fast localized spectral filtering. arXiv preprint arXiv:1606.09375 (2016)

5. Desikan, R.S., Ségonne, F., Fischl, B., Quinn, B.T., Dickerson, B.C., Blacker, D., Buckner, R.L., Dale, A.M., Maguire, R.P., Hyman, B.T., et al.: An automated labeling system for subdividing the human cerebral cortex on mri scans into gyral based regions of interest. Neuroimage 31(3), 968–980 (2006)

6. Fey, M., Lenssen, J.E.: Fast graph representation learning with PyTorch Geometric. In: ICLR Workshop on Representation Learning on Graphs and Manifolds (2019)

7. Frost, M.A., Goebel, R.: Measuring structural-functional correspondence: spatial variability of specialised brain regions after macro-anatomical alignment. Neuroimage 59(2), 1369–1381 (2012)

8. Gal, Y., Ghahramani, Z.: Dropout as a bayesian approximation: Representing model uncertainty in deep learning. In: international conference on machine learning. pp. 1050–1059. PMLR (2016)

9. Given, N.A.: Benchmarking geometric deep learning for cortical segmentation and neurodevelopmental phenotype prediction. In preparation (2021)

10. Glasser, M.F., Coalson, T.S., Robinson, E.C., Hacker, C.D., Harwell, J., Yacoub, E., Ugurbil, K., Andersson, J., Beckmann, C.F., Jenkinson, M., et al.: A multimodal parcellation of human cerebral cortex. Nature 536(7615), 171–178 (2016)

11. Glasser, M.F., Smith, S.M., Marcus, D.S., Andersson, J.L., Auerbach, E.J., Behrens, T.E., Coalson, T.S., Harms, M.P., Jenkinson, M., Moeller, S., et al.: The human connectome project’s neuroimaging approach. Nature neuroscience 19(9), 1175–1187 (2016)

12. Glasser, M.F., Sotiropoulos, S.N., Wilson, J.A., Coalson, T.S., Fischl, B., Andersson, J.L., Xu, J., Jbabdi, S., Webster, M., Polimeni, J.R., et al.: The minimal preprocessing pipelines for the human connectome project. Neuroimage 80, 105–124 (2013)

13. Glasser, M.F., Van Essen, D.C.: Mapping human cortical areas in vivo based on myelin content as revealed by t1-and t2-weighted mri. Journal of Neuroscience 31(32), 11597–11616 (2011)

14. Gordon, E.M., Laumann, T.O., Adeyemo, B., Gilmore, A.W., Nelson, S.M., Dosenbach, N.U., Petersen, S.E.: Individual-specific features of brain systems identified with resting state functional correlations. Neuroimage 146, 918–939 (2017)

15. Gratton, C., Kraus, B.T., Greene, D.J., Gordon, E.M., Laumann, T.O., Nelson, S.M., Dosenbach, N.U., Petersen, S.E.: Defining individual-specific functional neuroanatomy for precision psychiatry. Biological psychiatry (2019)

16. Havaei, M., Davy, A., Warde-Farley, D., Biard, A., Courville, A., Bengio, Y., Pal, C., Jodoin, P.M., Larochelle, H.: Brain tumor segmentation with deep neural networks. Medical image analysis 35, 18–31 (2017)

17. Heckemann, R.A., Hajnal, J.V., Aljabar, P., Rueckert, D., Hammers, A.: Automatic anatomical brain mri segmentation combining label propagation and decision fusion. NeuroImage 33(1), 115–126 (2006)

18. Kipf, T.N., Welling, M.: Semi-supervised classification with graph convolutional networks. arXiv preprint arXiv:1609.02907 (2016)

19. Kong, R., Li, J., Orban, C., Sabuncu, M.R., Liu, H., Schaefer, A., Sun, N., Zuo, X.N., Holmes, A.J., Eickhoff, S.B., et al.: Spatial topography of individual-specific cortical networks predicts human cognition, personality, and emotion. Cerebral cortex 29(6), 2533–2551 (2019)

20. Makropoulos, A., Robinson, E.C., Schuh, A., Wright, R., Fitzgibbon, S., Bozek, J., Counsell, S.J., Steinweg, J., Vecchiato, K., Passerat-Palmbach, J., et al.: The developing human connectome project: A minimal processing pipeline for neonatal cortical surface reconstruction. Neuroimage 173, 88–112 (2018)

21. Miller, K.L., Alfaro-Almagro, F., Bangerter, N.K., Thomas, D.L., Yacoub, E., Xu, J., Bartsch, A.J., Jbabdi, S., Sotiropoulos, S.N., Andersson, J.L., et al.: Multimodal population brain imaging in the uk biobank prospective epidemiological study. Nature neuroscience 19(11), 1523–1536 (2016)

22. Monti, F., Boscaini, D., Masci, J., Rodola, E., Svoboda, J., Bronstein, M.M.: Geometric deep learning on graphs and manifolds using mixture model cnns. In: Proceedings of the IEEE Conference on Computer Vision and Pattern Recognition (CVPR) (July 2017)

23. Ongür, D., Ferry, A.T., Price, J.L.: Architectonic subdivision of the human orbital and medial prefrontal cortex. Journal of Comparative Neurology 460(3), 425–449 (2003)

24. Reinke, A., Eisenmann, M., Tizabi, M.D., Sudre, C.H., Rädsch, T., Antonelli, M., Arbel, T., Bakas, S., Cardoso, M.J., Cheplygina, V., et al.: Common limitations of image processing metrics: A picture story. arXiv preprint arXiv:2104.05642 (2021)

25. Robinson, E.C., Garcia, K., Glasser, M.F., Chen, Z., Coalson, T.S., Makropoulos, A., Bozek, J., Wright, R., Schuh, A., Webster, M., et al.: Multimodal surface matching with higher-order smoothness constraints. Neuroimage 167, 453–465 (2018)

26. Robinson, E.C., Jbabdi, S., Glasser, M.F., Andersson, J., Burgess, G.C., Harms, M.P., Smith, S.M., Van Essen, D.C., Jenkinson, M.: Msm: a new flexible framework for multimodal surface matching. Neuroimage 100, 414–426 (2014)

27. Ronneberger, O., Fischer, P., Brox, T.: U-net: Convolutional networks for biomedical image segmentation. In: International Conference on Medical image computing and computer-assisted intervention. pp. 234–241. Springer (2015)

28. Sabuncu, M.R., Yeo, B.T., Van Leemput, K., Fischl, B., Golland, P.: A generative model for image segmentation based on label fusion. IEEE transactions on medical imaging 29(10), 1714–1729 (2010)

29. Sudre, C.H., Li, W., Vercauteren, T., Ourselin, S., Cardoso, M.J.: Generalised dice overlap as a deep learning loss function for highly unbalanced segmentations. In: Deep learning in medical image analysis and multimodal learning for clinical decision support, pp. 240–248. Springer (2017)

30. Van Essen, D.C., Glasser, M.F., Dierker, D.L., Harwell, J., Coalson, T.: Parcellations and hemispheric asymmetries of human cerebral cortex analyzed on surface-based atlases. Cerebral cortex 22(10), 2241–2262 (2012)

31. Van Essen, D.C., Smith, S.M., Barch, D.M., Behrens, T.E., Yacoub, E., Ugurbil, K., Consortium, W.M.H., et al.: The wu-minn human connectome project: an overview. Neuroimage 80, 62–79 (2013)

32. Zhao, F., Xia, S., Wu, Z., Duan, D., Wang, L., Lin, W., Gilmore, J.H., Shen, D., Li, G.: Spherical u-net on cortical surfaces: methods and applications. In: International Conference on Information Processing in Medical Imaging. pp. 855–866. Springer (2019)

